# mBER: Controllable *de novo* antibody design with million-scale experimental screening

**DOI:** 10.1101/2025.09.26.678877

**Authors:** Erik Swanson, Michael Nichols, Supriya Ravichandran, Pierce Ogden

**Affiliations:** Manifold Bio, Boston, MA, USA

## Abstract

Recent machine learning approaches have achieved high success rates in designing protein binders that demonstrate *in vitro* binding to their targets. While open models for unconstrained “minibinder” design have shown great experimental promise, methods for designing binders in specific formats, such as antibodies, have lagged behind in experimental success rates. In this work, we present mBER, an open-source protein binder design system capable of designing antibody-format binders with state-of-the-art experimental success rates. mBER builds on the ColabDesign framework, achieving successful antibody design primarily through the inclusion of informative sequence and structure conditioning information. Using mBER, we designed two libraries comprising over 1 million VHH binders against 436 diverse targets. We experimentally screened the two libraries against 145 of these targets, resulting in a dataset of over 100 million binding interactions. We achieved specific and significant design success against 45% of targets. In a filtered set of designs, we detect binding rates to specific epitopes as high as 38%. Through mBER, we demonstrate that format-specific binder design is possible with no additional training of underlying folding and language models. This work represents the largest reported *de novo* protein design and validation campaign, and one of the first open-source methods to demonstrate double-digit percentage experimental success rates for antibody binder design.

## 1 Introduction

In the development of protein therapeutics, discovery of binders with specific affinity against targets of interest is a critical step. It has long been a goal of protein engineering to achieve rational design of such binders, enabling fine-grained control over biophysical properties relevant to developability and clinical efficacy [1].

Traditional methods for binder discovery often involve large-scale *in vitro* screening of naive or immunization-derived libraries [2, 3, 4]. These methods produce binders with little control over sequence composition, epitope, or paratope, all of which can determine the success or failure of a drug in the clinic. Performing a series of follow-up assays to determine and filter on these properties is costly and time-consuming and offers no guarantee of reaching an optimal candidate from an initial set of binders.

Recent advances in AI for molecular modeling have opened up new avenues of binder discovery, enabling *de novo* design of structure and sequence for epitope-targeted binders. To date, diffusion- and loss optimization-based methods have demonstrated the ability to design binders against diverse target sets with high experimental hit rates. Popular design methods such as RFDiffusion [5] and BindCraft [6] excel at generating unconstrained “minibinders”, small proteins with completely novel structures and sequences.

While these methods offer control over epitope targeting and some control over binder geometry, they fall short in their ability to impose constraints on binder sequence and structure. These design constraints are necessary for format-specific protein design, specifically the design of antibody-format drugs, which contain large, highly conserved framework regions, as well as structured diversity within variable regions [4, 7].

Antibodies represent the dominant format for protein therapeutics, with over 100 FDA-approved antibody drugs and established manufacturing, regulatory, and clinical development pathways. Decades of clinical experience have established predictable pharmacokinetics, immunogenicity profiles, and safety characteristics for antibody scaffolds [8]. Diverse antibody modalities such as single-domain VHHs are popular therapeutic formats for their small size and developability [9, 10]. The ability to design antibodies thus represents a more immediately translatable advance relative to minibinder design, combining the control and epitope-specificity of structural design with the proven therapeutic utility of the antibody format.

Multiple groups have reported success in generating *de novo* antibodies with some *in vitro* experimental success, including JAM and Chai-2 [11, 12]. To date, RFAntibody [13] and Germinal [14] are the only open-source models with experimental evidence supporting claims of successful *de novo* antibody design. Germinal appeared contemporaneously with this work and presents similar methods and experimental success rates to our own; RFAntibody, in contrast, remains more computationally intensive with lower yields.

In this work, we present Manifold Binder Engineering and Refinement (mBER), a protein binder design system building on the work of ColabDesign and BindCraft that enables design of antibody binders with partially constrained sequences and structures. As our core contribution, we provide structural templates and sequence priors for VHH-format binders to AlphaFold-Multimer, and show that this conditioning is sufficient to guide backpropagation-based binder design of high-confidence multimeric folded structures. Using mBER, we design a pair of libraries totaling more than 1 million VHH-format binders to a set of 436 diverse targets, and experimentally screen these libraries against 145 of these targets, resulting in a rich dataset of on- and off-target binding interactions. Our libraries achieve specific and significant success against 65 of these targets, with per-binder success rates as high as 38% for some epitopes. We propose mBER as a method to generalize protein design to novel formats without requiring additional model training or finetuning to formats of interest.

## 2 Results

### Building on AlphaFold2 Backpropagation Design Methods

The experimental success of minibinders generated by BindCraft [6] has drawn attention towards backpropagation-based protein design methods. BindCraft is based on ColabDesign, a framework with utilities for preparing inputs to AlphaFold2, computing losses on outputs from the model, and backpropagating gradients from those losses to the input space for continuous and discrete sequence design. Prior to BindCraft, several approaches leveraged ColabDesign or related AlphaFold-based methods for protein design, but until very recently none had achieved experimentally validated antibody design [15, 16, 17, 18, 19]. Concurrently with our work, Germinal [14] independently reported a similar backpropagation-based approach with experimental success, underscoring the rapid emergence of this design paradigm.

Given the general success of AlphaFold-based protein design methods, we reasoned that antibody design may be possible by incorporating informative structure and sequence conditioning at design time. To this end, we developed mBER—a versatile framework for creating binders that integrates sequence and structural constraints to achieve precise control over the resulting proteins.

In Figure 1, we show a schematic of the mBER design system for VHH. As inputs, a target structure with optional hotspots is provided, alongside a partially-masked VHH framework sequence. mBER prepares an informative structural template and sequence prior from these inputs and uses them to produce high-quality binder designs via backpropagation through AlphaFold-Multimer [20]. The final designs are then re-folded and scored with *in silico* confidence metrics. We test designs against targets in an all-against-all fashion, obtaining a rich matrix of on-target and off-target binding data.

**Figure 1:**
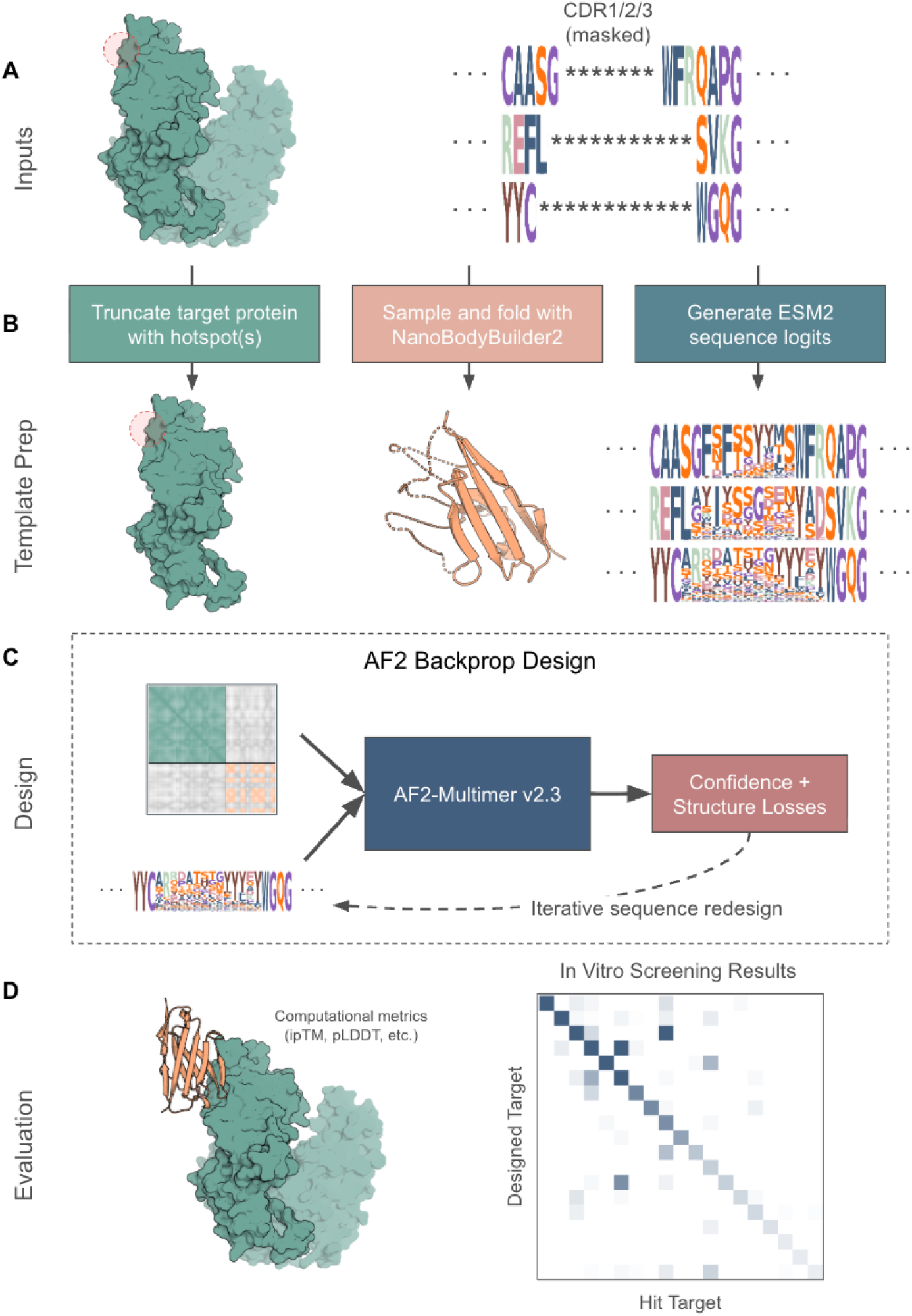
Schematic Overview of the mBER design system for VHH. **(A)** An input state consisting of a target structure (with optional hotspot) and a partially masked framework sequence are provided by the user. **(B)** mBER performs automated truncation of target proteins to a smaller, easily foldable epitope. The input masked sequence is passed through ESM2 to generate a probability distribution over the masked tokens in the form of sequence logits. A single sequence is sampled and folded with NanoBodyBuilder2 to produce a structural VHH template. **(C)** Template features are provided, along with a continuous sequence representation, to AlphaFold-Multimer. We compute losses based on AF2’s confidence module and predicted structure. The input sequence is iteratively optimized with respect to these losses. **(D)** A final forward pass through AlphaFold-Multimer yields a folded structure and confidence for each generated sequence. We experimentally validate our designs across a large set of targets.

### Experimental Validation of More Than 1 million VHH Binder Designs

We selected 436 human cell-surface proteins as candidate targets for tissue-specific drug delivery vehicles. Across two large-scale phage-display experiments, we designed 1,153,241 unique VHH binders against this target set. Of the 145 targets evaluated experimentally, 65 (45%) exhibited significantly elevated on-design hit rates under Fisher’s exact test (Figure 2B; Supplementary Information A.11). These results demonstrate that *de novo* computational design can reliably generate VHH binders to a substantial fraction of diverse targets in a multiplexed setting.

**Figure 2:**
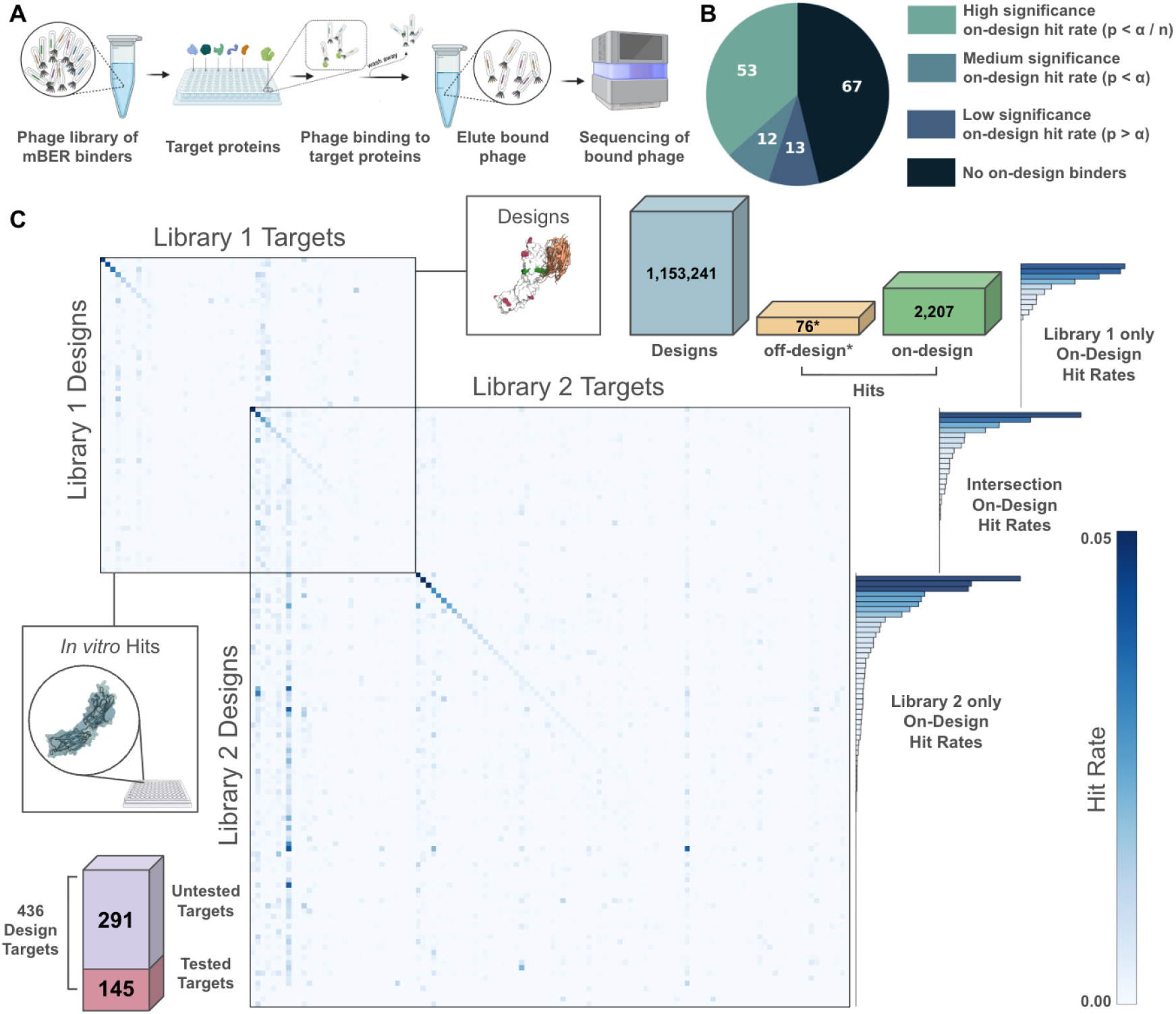
Large-scale validation of over 1 million designed VHH binders. **(A)** Schematic of the multiplexed phage screening pipeline: designed VHH sequences are displayed on phage, panned against immobilized targets, and sequenced by NGS to assess binding. **(B)** Statistical outcomes across 145 tested targets: pie chart summarizes significance levels of on-design binding relative to off-design background (significance level *α* = 0.05; Bonferroni correction *n* = 145). **(C)** All-against-all hit rate matrix for designs and targets in both experiments. Strong diagonal enrichment highlights targets where large fractions of designed binders produced on-design hits. Bars show on-design hit rates for targets with any on-design hits, split by experiment and by intersection. ^*^Off-design hit count is normalized by target count.

Our validation pipeline (Figure 2A) leverages a phage-based, all-against-all binding assay: each library of designed VHHs is expressed on phage, panned against immobilized target proteins in 96-well format, and sequenced by NGS to recover the identities of bound binders. This yields comprehensive readouts of binding signal across both intended and off-targets. To interpret these data, we apply a stringent order-statistics–based hit-calling criterion (Supplementary Information A.10), which ensures hits are detectable, consistent across replicates, and specific relative to background and off-target binding. Parameters are chosen such that the expected number of false positives per target is less than one across the entire library.

The two phage experiments provide complementary views of mBER performance. In the first, *Library 1*, we generated 565,982 binders against 370 targets and screened them against 61 proteins, yielding hits to 56 targets, including 22 (36%) with significantly elevated on-design hit rates. Building on this success, we expanded to a second library, *Library 2*, comprising 587,259 binders against 389 targets, screened against 116 proteins. Here, 115 of 116 targets yielded at least one hit, and 49 (42%) showed significantly elevated on-design hit rates. Together, these experiments underpin the robustness of our binder design and screening pipeline.

Due to the all-against-all nature of our binding assay, we observe a large number of binding interactions between binders and targets which they were not explicitly designed against. We designate hits where the designed target matches the hit target “on-design” and hits where the designed target does not match the hit target “off-design”. We distinguish this notion from off-target binding, as both on- and off-design hits are typically specific in signal to their hit targets (Supplementary Figure S1).

From 1,153,241 designs, we observe 36,008 off-design hits, many of which arise from a handful of seemingly “sticky” targets, which appear as column-like striations in a heatmap of design targets against hit targets (Figure 2C). A subset of 331,578 designs had targets in the experimental set, accounting for 11,040 off-design hits and 2,207 on-design hits. While the large number of off-design hits is at first surprising, we reconcile it by noting that the probability of discovering an off-design binding interaction increases dramatically with the number of targets tested. As on-design hits can only be called to a single target, and off-design hits can be called to 145, we normalize the 11,040 by the number of targets and note that the appropriate comparison is about 76 off-design hits to 2,207 on-design hits.

### Per-target hit rates under ipTM filtering

The principal distinction between the *Library 1* and *Library 2* libraries lies in their ipTM distributions. AlphaFold-Multimer’s ipTM metric has previously been shown to correlate with binding success when used as a filter for *de novo* binder designs [6, 5, 13, 21]. To assess the utility of ipTM in our multiplexed binding assays, we systematically examined how these metrics tracked with experimental outcomes.

78 of the 145 experimentally tested targets showed some degree of on-design binding, with a wide variation in hit rates per designed binder. These hit rates range between approximately 0.02% and 8%, with a median hit rate of approximately 0.4%. Applying successive ipTM filters substantially increases effective hit rates on a per-binder and per-target basis, with hit rates in the double-digit percent range when filtering to designs with ipTM scores greater than 0.7, and a majority of hit rates surpassing 1% under an ipTM filter of 0.8 (Figure 3A).

**Figure 3:**
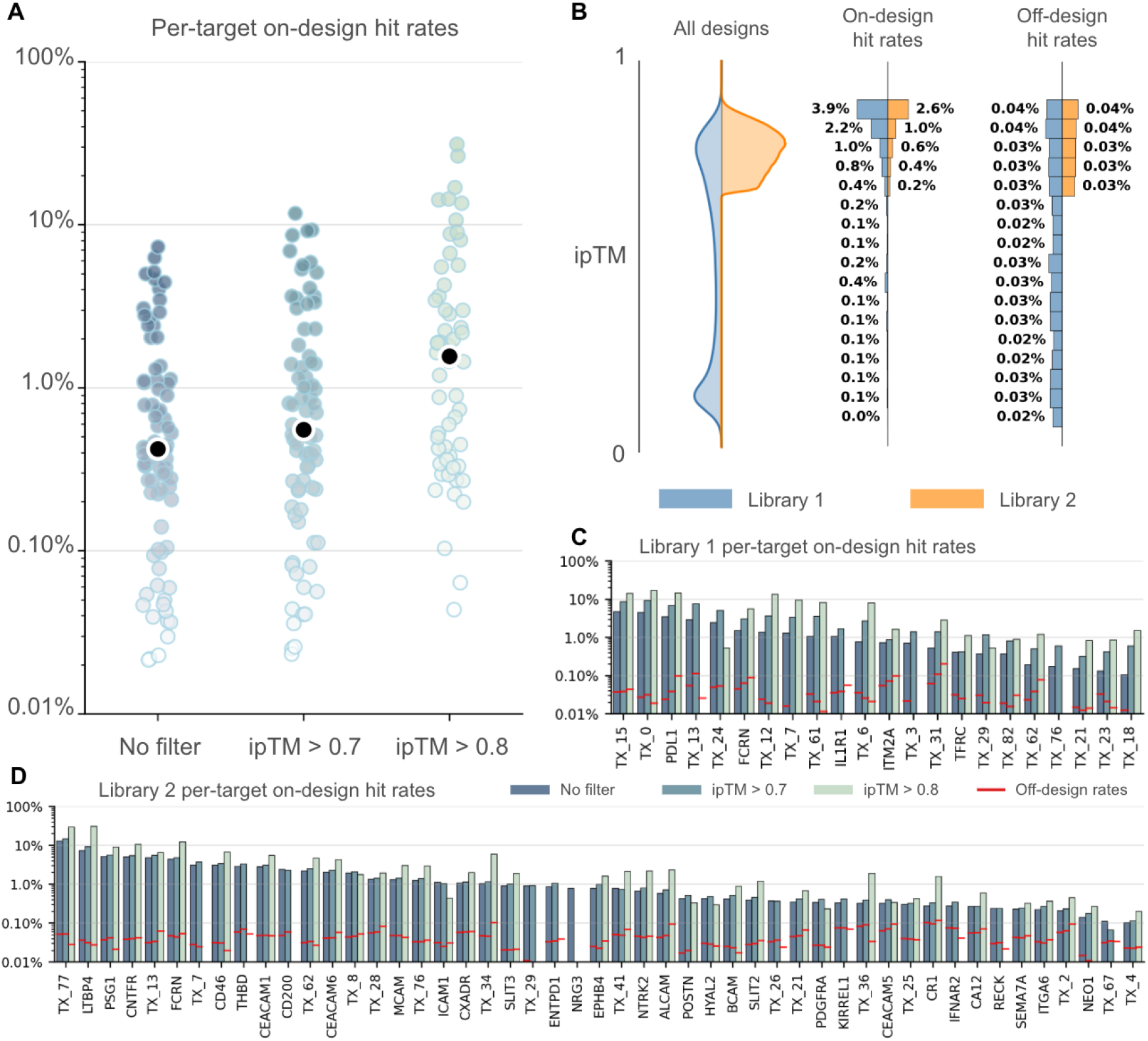
AlphaFold-Multimer metrics as effective filters for extracting on-design hits. **(A)** Effective on-design hit rates for 78 targets under different filtering criteria (67 targets with no designed hits are not shown). The median hit rate in each set is shown in black. **(B)** Distribution of on- and off-design hit rates binned by ipTM cutoffs of width 0.05. We filter out bins containing less than 0.1% of each library. Off-design hit rates are normalized by the number of unique targets tested in each library experiment. **(C, D)** Effective on-design hit rates for all targets with significantly elevated on-design hit rates under different filtering criteria. Off-design hit rates are shown in red. Proprietary targets are anonymized with names of the form `TX_N`.

In *Library 1*, the ipTM distribution closely followed that of naive mBER generations, with a bimodal profile: one population of low-confidence binders centered around 0.15 and another of high-confidence binders centered around 0.8. By intentionally sampling across this range, we confirmed that experimental hit rates rise sharply at higher ipTM values, reaching 3.9% for binders in the 0.85–0.9 range. Importantly, off-design hit rates remained relatively flat across ipTM bins, supporting the conclusion that ipTM filtering enriches for functional binders without elevating polyspecific binding (Figure 3B).

Building on these findings, we constructed *Library 2* by restricting to designs with ipTM > 0.66, yielding a unimodal high-confidence distribution. Although absolute hit rates in the top ipTM bins were slightly reduced relative to *Library 1*, ipTM remained a strong predictor of success. In both libraries, we also investigated pLDDT (averaged across all residues) as a predictor of on- and off-design binding. While pLDDT performed comparably or worse than ipTM as a predictor of on-design hits, there appears to be a weak correlation between pLDDT scores and off-design binding (Supplementary Figure S2). We speculate that pLDDT, being largely reflective of monomeric fold quality, tends to favor generally well-structured VHHs, which may display higher baseline binding propensity regardless of the intended target.

Finally, we evaluated ipTM filtering on a per-target basis. Applying thresholds of ipTM > 0.7 and > 0.8 consistently increased effective hit rates, in some cases by an order of magnitude relative to the unfiltered libraries. In *Library 2*, ipTM scores for most binders already exceeded 0.7, resulting in minimal change at that cutoff, but applying the 0.8 filter boosted effective success rates for several targets. Despite this enrichment, the overall library-wide on-design hit rate increased only modestly from 0.5% in *Library 1* to 0.7% in *Library 2*. Nonetheless, for individual targets such as TX_77 and LTBP4, strict filtering produced hit rates exceeding 30% (Figure 3C,D).

### mBER generates binders to multiple epitopes with differential success rates

During design of both *Library 1* and *Library 2*, we selected hotspots randomly from highly exposed surface residues on the target protein structures and truncated the target at design time to a set of residues containing the selected hotspot (Supplementary Information A.4). This hotspot selection strategy was chosen to maximize the number of unique epitopes targeted by mBER designs. We evaluated binders with both high and low *in silico* confidence scores across these hotspot choices to assess mBER’s ability to achieve epitope-specific success.

Generally, we find that experimental success with mBER designs is highly sensitive to the choice of epitope, with on-design hits to only a few hotspots per target. For some targets, such as TFRC (Figure 4A), we observed binders to multiple epitopes, but low *in silico* scores and small sample sizes limit confidence in these hits. By contrast, for FCRN (Figure 4) we recovered a large number of high-confidence binders spanning three distinct epitopes, and in one hotspot we detected multiple distinct binding geometries—demonstrating mBER’s capacity to diversify paratopes as well as epitopes. In other cases, hotspotting did not fully constrain binder positioning. In the case of PDL1 (Figure 4C), nearly all successful binders clustered to a single epitope and geometry despite designs targeting multiple hotspots, including those positioned on the opposite side of the domain.

**Figure 4:**
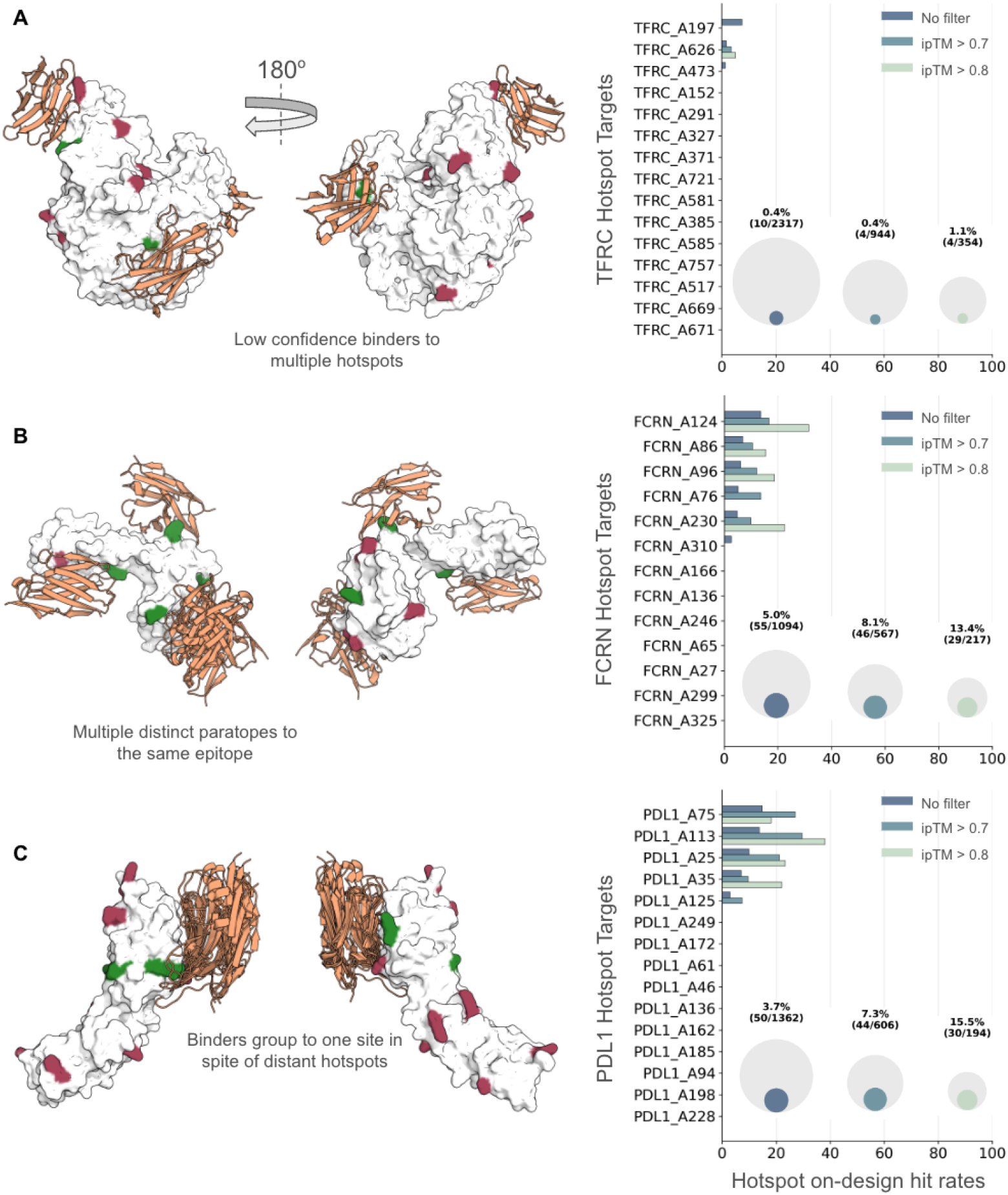
Epitope-specific behavior of mBER. A subset of de-anonymized target structures are shown. Hotspots with any number of experimentally successful designs are shown in green, while hotspots with no successes are shown in red. For each successful hotspot, a single binder to that hotspot is shown in orange. Experimental success rates for designs against each hotspot are shown to the right, under successive ipTM filters. The total binder success rates to these targets are also shown as nested circles, where the outer circle represents the total number of designs, and the inner circle represents the total number of experimental hits. **(A)** Against TFRC, we achieve experimental success for a few binders to 3 distinct epitopes. However, the ipTM scores for some binders are low, and the total number of successful binders in this set is small. **(B)** Against FCRN, we call hits from binders designed against 3 unique epitopes, with high ipTM score. For one epitope, multiple distinct paratopes are visible. **(C)** PDL1 shows high design success rates, close to 40% for the best hotspot under strict ipTM filtering. However, all the binders group to a single site, even when designed against hotspots on the opposite side of the targeted domain.

These results highlight the strong epitope dependence of design outcomes. Because each hotspot functions effectively as a distinct target, success rates are substantially higher when measured at the epitope level compared to the aggregate target level reported in Figure 2. For instance, the overall success rate for TFRC without filtering was only 0.4% (10/2317), yet its best-performing hotspot achieved 7.5% (6/80). Under strict ipTM filtering, highly favorable epitopes reached success rates as high as 38% (8/21), underscoring the importance of epitope selection in determining design efficiency.

## 3 Discussion

In this work, we have presented mBER, an open-source protein binder design system that leverages backpropagation through AlphaFold-Multimer to enable highly controllable, format-specific binder design. We employed mBER to design 1,153,241 binders against 436 human cell-surface protein targets using computational predictions of their structures. In our experimental validation, we raised thousands of specific binders with prospective epitope information against 78 of 145 tested targets, 65 of which display statistically elevated on-design hit rates. These binders form a large candidate set of tissue-targeted delivery vehicles for further exploration *in vivo*.

Our results validate ipTM as an effective computational filter for design selection, achieving success rates up to 38% for the most favorable epitopes after strict ipTM filtering. This finding supports the use of AlphaFold-Multimer’s confidence metrics as predictive indicators of design quality. These experimental success rates are comparable to state-of-the-art closed-source models such as JAM and Chai-2 and the open source method Germinal [11, 12, 14]. Our approach demonstrates that successful antibody-format binder design can be achieved without requiring additional training or fine-tuning of underlying folding and language models.

A key innovation of our approach is the effective combination of protein language model priors with structural templating to guide AlphaFold-Multimer toward generating functional VHH binders. By using protein language models to generate sequence distributions for masked CDR regions while maintaining fixed framework residues, we successfully balance the need for sequence diversity in binding regions with the structural constraints required for proper antibody folding. The addition of partial structural templates further stabilizes the design process, enabling AlphaFold-Multimer to produce high-confidence folds in the absence of MSA information.

### Limitations

Successful designs with mBER are limited to a subset of epitopes on a subset of targets. This constraint is shared across existing binder design methods, which remain unable to reliably generate binders to all epitopes on all targets. This introduces both practical and translational challenges. From a practical perspective, large amounts of computational and experimental effort are invested in epitopes that yield no productive binders. From a translational perspective, many therapeutic applications demand targeting of specific epitopes to block defined protein–protein interactions or to avoid competition with endogenous ligands. Overcoming these limitations will require mechanistic insight into why current methods fail on particular epitopes, enabling more effective epitope selection and informing the development of improved design algorithms.

Computational cost also represents a key limitation. While deeper sampling can improve *in silico* metrics such as ipTM and pLDDT that correlate with experimental success, this approach becomes prohibitively expensive at scale. Similar trends have been observed in other generative binder design methods, where increased test-time computation enhances accuracy [22]. These findings highlight the importance of reducing computational demands and improving sampling efficiency, which could yield higher design success rates without increasing overall cost.

Our library-based screening approach offers both notable strengths and clear limitations. It enables us to evaluate binding for an unprecedented number of de novo designs against hundreds of targets in an all-versus-all format. This dataset, comprising over 100 million protein–protein interactions, provides rare insight into off-design binding, an underexplored but important consideration in binder engineering. Our large target set allows us to directly assess and avoid non-specific binding instead of relying on polyreactivity reagents. The statistical enrichment of on-design over off-design hits strongly supports epitope-specific engagement; however, we cannot confirm the predicted binding modes without structural validation, and more direct experimental measurements will be necessary to verify agreement between designed and actual binding conformations. In addition, our phage-display assays do not permit precise quantification of binding affinities. We expect affinities for our binders lie primarily in the micromolar to nanomolar range, as millimolar interactions would not survive selection [23]. The specificity observed in our binders supports the view that very weak interactions are not captured by our screen, since these would be expected to appear as widespread cross-reactivity across targets.

### Future Directions

The mBER framework opens several avenues for future research and development. mBER’s loss functions and hyperparameters are largely inherited from ColabDesign and BindCraft with few modifications beyond mBER’s structure and sequence templating. This hyperparameter space has not been exhaustively explored, leaving open the possibility that significant improvements could be achieved by refinement. Key parameters including loss function weights, optimization schedules, hotspot selection, and target truncation could all impact design success. Moreover, optimal parameters likely vary by target, epitope, and binder format, suggesting that adaptive hyperparameter selection could improve performance.

Beyond hyperparameter optimization, the control mBER affords over binder format presents opportunities for expanding the designable epitope space. Different epitope topologies may be better addressed by different binder formats. mBER’s templating and masking approach can be applied to any protein scaffold by supplying appropriate structural templates and sequence priors. Extending design beyond VHHs may therefore broaden the spectrum of accessible epitopes and targets. Larger antibody formats such as scFv or Fab fragments could provide additional paratope surface area, while non-antibody scaffolds like designed ankyrin repeat proteins (DARPins), fibronectin domains, or novel formats invented with computational designability in mind may each offer distinct geometric and biophysical properties [24, 25]. mBER’s format-controlled approach enables matching scaffold characteristics to epitope topology, an underexplored dimension of rational binder design.

Because backpropagation-based design methods depend on the accuracy of the underlying protein folding model, advances in folding models can be expected to directly improve design outcomes. AlphaFold2’s folding accuracy has been surpassed by models such as AlphaFold3 and Boltz-2, particularly in the domain of antibody folding [26, 27]. Molecular design by backpropagation on these models has been explored [28]; however, both AlphaFold3 and Boltz-2 utilize diffusion-based structure modules that complicate gradient propagation from predicted structures back to input sequences. The highly controllable and successful backpropagation process demonstrated by mBER may motivate training of improved protein folding models with gradient-based design applications in mind.

### Conclusion

mBER represents a significant advance in controllable protein binder design, demonstrating that format-specific binders can be successfully designed without requiring specialized model training. By combining the strengths of protein language models, structural templates, and AlphaFold2’s folding and confidence capabilities, we provide a flexible framework that can be adapted to various binder formats and design constraints. As the field continues to evolve with improved folding models and design strategies, approaches like mBER that leverage existing models in novel ways will be crucial for democratizing access to protein design capabilities and accelerating therapeutic discovery.

## Acknowledgements

We would like to thank the entire team at Manifold Bio for their research and operational contributions which have enabled and supported this work. Specifically, we thank Marshall Case and Alex Reis for many helpful discussions around the development of mBER and Sarah Post for assisting with the experiments reported in this paper. We thank Jeffrey Chang, Laksh Aithani, Kalyan Palepu, Rohil Badkundri, and Avery Parr for early reviews of the manuscript and Guocong Song for early reviews of the open-source codebase. We would also like to thank Sergey Ovchinnikov, Martin Pacesa, and the Bruno Correia lab for their contributions to open-source protein design tools, specifically ColabDesign and BindCraft. We are grateful for the support of AWS in providing computational resources for this work.

## Author Contributions

E.S., M.N., and P.O. conceived the work. E.S. and M.N. wrote the manuscript, which was edited by all authors. Analysis and code development were performed by E.S., with support from M.N. and P.O. E.S. and M.N. ran design campaigns and selected designs for experiments. S.R. performed all experimental work, including protein sourcing, library preparation, binding assay, and sequencing.

## Code Availability

Open-source code for running mBER is made available under an MIT license at https://github.com/manifoldbio/mber-open.

## A Supplementary Information

### A.1 mBER *in silico* VHH design protocol

The core design workflow of mBER consists of 4 distinct steps: input preparation, template preparation, design, and evaluation. In the mBER codebase, we provide a set of default configurations and a default design pipeline so that users may run VHH design in an identical fashion to how we designed the libraries reported here.

An mBER VHH design run is initialized from a target UniProt ID [29], RCSB PDB ID [30], or custom .pdb filepath. For UniProt IDs, a structure is downloaded from the AlphaFold Protein Structure Database [31]. In addition to the target structure, users can optionally provide strings indicating subregions and hotspots of the target, which will be used in the truncation process and during design to guide binders towards specific epitopes. A partially-masked VHH sequence is also required to initialize a design run. For the libraries reported in this paper, we use a set of VHH frameworks based on the human IGHV3-23 germline framework, which is commonly used for VHH humanization campaigns [32]. As our frameworks differ by only a few mutations from the human germline sequence, we expect a low risk of immunogenicity for our designed VHH binders. We provide a table of VHH frameworks and their comparison to IGHV3-23 in Table S1.

During template preparation, the target protein structure is truncated based on provided region and hotspot configurations. In the VHH protocol, ESM2 is used to generate a set of logits over masked regions of the VHH framework, which are converted to position-wise probabilities under a softmax operation. From these probabilities, we randomly sample a single sequence and produce a folded structure using NanoBodyBuilder2. The folded structure is combined with the truncated target structure into a single .pdb file and passed along with the ESM2 sequence logits to the design stage.

The design stage follows the `design_3stage` method in ColabDesign, with modifications from the binder protocol to enable separate templating of target and binder structures, while masking inter-chain contact maps. Sequence bias is incorporated as a logits vector which is added to the design logits prior to each design step. The sequence vector,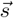, used to prepare inputs at each step for AlphaFold-Multimer can be written as

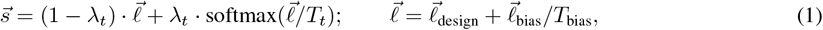

where λ_*t*_ is a schedule parameter which increases over design steps, *T*_*t*_ is a temperature parameter which decreases over design steps 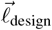 is the logits vector which is directly updated in the backpropagation design process, 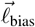 is the sequence bias logits vector generated by ESM2, and *T*_bias_ is a temperature parameter to scale the effect of sequence bias. As in BindCraft [6], we optimize primarily on the predicted aligned error (pAE) matrix across our binder and target chains, with auxiliary loss functions based on other AF2 confidence metrics, residue contacts, and sequence features. We use weights from 4 of 5 pre-trained AlphaFold-Multimer models for design, saving the last weight set for the evaluation stage. We find that providing antibody sequence bias and partial templates is sufficient to direct ColabDesign towards the design of functional VHH binders.

After the design stage, candidate binder sequences are passed along with the initial binder and target templates to an AlphaFold-Multimer model, initialized with the unused set of weights. Other than the weight set, our evaluation model is prepared with the same configuration as our design model, including template information for the binder and target structures. For both the design and evaluation stage, we use 3 recycles.

### A.2 Antibody Templates Guide AlphaFold Towards Successful VHH Binder Generation

Empirically, we have found that AlphaFold-Multimer, which struggles to accurately predict antibody-antigen interactions [20, 26], can be biased towards high-confidence folds by supplying an informative template, without any MSA information. In particular, templating enables the generation of designs with high ipTM scores, which have been shown to correlate well with binding for *de novo* designs [13, 6].

In mBER, we use templates to guide folding of both target and binder proteins. For target proteins, we select hotspots and truncate to a subset of target residues in order to improve design speed and targeting. We describe the truncation process in Supplementary Information A.4. For binders, we provide a single template in the form of a reference structure, matching the desired binder framework. For VHH, we typically generate the reference structure using the VHH-specific folding model NanoBodyBuilder2 [33], which runs quickly and does not rely on MSA information. Generating reference structures on the fly enables the use of arbitrary VHH frameworks, for maximal flexibility of design constraints. We combine the VHH and the truncated target structures into a single .pdb file and provide them as templates to AlphaFold-Multimer during both design and evaluation. We mask inter-chain information, as well as template information in CDR loops, to allow for flexible binding.

In Figure S4A, we show the effect of providing target and VHH templates at design time, initializing the design sequence as a uniform distribution over amino acids in every position. We find that templating alone is insufficient to steer AlphaFold-Multimer towards effective VHH design, and does not strongly constrain the secondary structure of the predicted binder. Better sequence priors are required to guide successful binder generation.

### A.3 Protein Language Models Provide Flexible Sequence Priors for Antibody Binder Formats

A major challenge in *de novo* antibody design is the application of sequence constraints. Antibody variable chains are made up of fixed framework sequences interspersed with highly variable complementarity determining regions (CDRs) [4, 7]. In general, the antibody design problem requires one to fix certain framework residues, while designing realistic CDRs which fold well in the context of their frameworks. Previous work has shown that common structure-to-sequence design tools like ESM-IF and ProteinMPNN fail to capture the natural distribution of amino acids in antibody CDRs [34], but that such methods can be rescued by ensembling with sequence-to-sequence predictions from protein language models (PLMs) [35].

In mBER, we use ESM2 [36] to provide sequence guidance during binder design. As inputs to our VHH design protocol, a partially-masked VHH framework sequence is provided. Typically, we choose to mask all 3 CDR regions, while fixing framework residues, although mBER can also accommodate masking of additional framework positions and fixing of CDR residues. We run a single forward pass of the partially masked sequence through a chosen PLM and extract the logits over the masked positions. After scaling by some temperature, these logits are converted to position-wise probabilities, resulting in a *L* × 20 vector, where *L* is the length of the full VHH sequence and each of 20 dimensions represents the probability of the corresponding amino acid occupying that position. The sequence logos in Figure 1B represent the positional probability distributions over amino acids in a set of CDRs. The resulting sequence vector is used to initialize the ColabDesign optimization process. The logits are further used as a bias term to steer sequence design across all optimization steps. We describe the sequence guidance process in more detail in A.1. In our code, we also provide the option to use AbLang2 [37] for sequence guidance.

In Figure S4B, we show the effect of providing a VHH sequence prior during design, while providing structural templates only for the target protein. While the resulting design is somewhat VHH-like in structure, it contains more disordered loops than a typical VHH and poor local confidence scores, as measured by pLDDT. In Figure S4(C), we show that providing both a VHH sequence prior and a VHH template during design leads to high confidence folds, with plausible docked VHH structures.

### A.4 Epitope Selection and Target Truncation

Computationally predicted structures from the AlphaFold Protein Structure Database (AFDB) were obtained for each target [31]. Target proteins were first truncated to their extracellular domains as identified by TMbed, a protein language model predictor of transmembrane domains [38]. Next, surface exposed residues were identified by their solvent-accessible surface areas (SASA) in the predicted structures. In each mBER design trajectory a random hotspot residue was chosen among these surface exposed residues, leading to at least 10 hotspot residues sampled per target. To reduce computational cost the target protein was then truncated using a dynamic programming approach to omit residues beyond 25 angstroms from the hotspot while minimizing chain breaks in the resulting structure. This truncated structure centered on the hotspot was then provided to mBER as the design target.

### A.5 *In Silico* Comparison to RFAntibody Designs

We compare mBER to the open source antibody design method RFAntibody. In an attempt to run a fair comparison, we provide exactly the same inputs to mBER and RFAntibody, using the default parameters in the RFAntibody source code. Specifically, we provide a truncated target structure and a VHH scaffold. We generate one diffusion trajectory per input config and sample 10 ProteinMPNN sequences from the resulting complex structure. With mBER, we similarly run one trajectory and sample 10 mutagenesis sequences for each input config.

All resulting binders are evaluated under the AF2 pipeline used in mBER, where structural templates for both the binder and target (but not their interactions) are provided to AlphaFold-Multimer to refold with 3 recycles. We collect and report ipTMs gathered from these evaluation folds in Figure S3. We find that mBER consistently produces binders with higher ipTM scores under our *in silico* evaluations. A limitation of RFAntibody is that many generations are often required to produce a single high-quality design [13]. In practice, we find that it is *not* necessary to generate many mBER binders before finding a single *in silico* hit, so long as the inputs are reasonable.

### A.6 Protein Sourcing

All proteins were either produced in-house or sourced from Acro Biosystems. For in-house production, protein sequences were obtained from Uniprot [29] and truncated to extracellular domains as described in Supplementary Information A.4. Sequences for resulting extracellular domains were codon optimized for mammalian expression and ordered as gene blocks with a capture tag (eg:6xHis) on the C-terminus for ease of capture. Commercially sourced proteins included Human PDL1, Cat. PD1-H5229; Human FcRn, Cat. FCN-H52W; Human TfRC, Cat. CD1-H5243 (ACROBiosystems).

Proteins were produced in-house using the Expi293 HEK-based expression system according to manufacturer instructions. Cells were pelleted by centrifugation and proteins were purified from supernatants using a magnetic Protein A bead based system. Purified molecules were quantified using a Bicinchoninic Acid (BCA) assay and construct sizes were confirmed by SDS-page gel. Proteins that successfully expressed were desalted into PBS using IMCS SizeX Tips on a Hamilton liquid handler and an equimolar amount of each protein was aliquoted, flash-frozen with liquid nitrogen, and stored at -80°C until further use.

### A.7 Library Cloning

Libraries were cloned into VHH frameworks using standard Gibson Assembly protocols. Library inserts and the vector were combined at optimized ratios with assembly master mix and incubated under standard conditions. The assembled fragments were electroporated into bacterial cells and transformed colonies were analyzed for library coverage. Once adequate coverage was confirmed, libraries were prepared and stored at -80°C until further use.

### A.8 Phage Production

Bacterial cells for phage production were grown in supplemented media at 37°C with shaking until reaching optimal density. Cells were infected with helper phage and incubated for 1 hour. The infected cells were then resuspended in media containing antibiotics and incubated overnight with gentle shaking. Phage were PEG precipitated and purified. Once purified, phage were titered in bacterial cells, and stored at 4°C until use.

### A.9 Phage Display for Binding Evaluation

Proteins were coated onto appropriate capture plates. Plates were washed and incubated with a blocking solution for 1.5 hours at room temperature. Phage libraries at appropriate titers were added and incubated for 2 hours. After washing to remove unbound phage, bound phage were eluted enzymatically and used to infect bacterial cells. Infected cells were selected overnight in antibiotic-containing media with shaking. Samples were prepared for next-generation sequencing using appropriate PCR primers to amplify binder sequences, then uniquely indexed and sequenced.

### A.10 Calling Hits from Phage Data

We call hits on a per-sequence basis using order statistics over the signal measured to that sequence in every well of our assay. In order to ensure that signal across different wells is comparable, in addition to the phage library, we add a spike in control (SIC) sequence at a known and fixed concentration to every well. The SIC provides a reference NGS counts signal to which we can normalize our counts for every other sequence in the library. Specifically, for a given sequence *i* and well *j*, we compute a normalized signal *ñ*_*i,j*_ from the raw counts *n*_*i,j*_ and the SIC counts *n*_SIC,*j*_ as

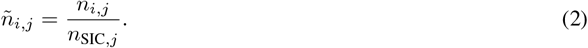

For a target *k*, we then consider the set *K* of indices corresponding to wells where that target is immobilized and compute a hit-calling ratio *r*_*i,k*_ as

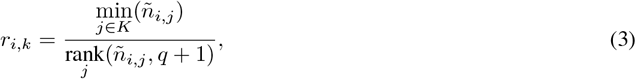

where 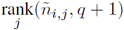 corresponds to the (*q* + 1)-th largest element *ñ*_*i,j*_, taken over the index *j*. We call sequence *i* a hit to target *k* if *r*_*i,k*_ > 1.

To establish a false positive rate for this hit-calling method, we consider the case of a sequence s which does not bind specifically to any target. In such a case, we expect that *ñ*_*s,j*_ ∼ i.i.d over wells *j*. In other words, the signal we expect from such a sequence should be drawn from the same distribution in every well. We can apply order statistics to remain distribution free and estimate the probability that *r*_*s,k*_ > 1 for a given target *k*.

*r*_*s,k*_ > 1 is equivalent to the statement that all wells in the set *K* have higher signal to the sequence s than the (*q* + 1)-th highest signal well 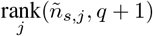 (this can only occur if *q* > |*K*|). Supposing that there are *N*_*wells*_ total wells, there are 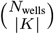 unique sets of ranks for the *K* wells and 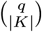 unique sets of ranks where all have higher signal than the (*q* + 1)-th well. Under the i.i.d assumption, all possible rankings are equally likely, and thus the probability of calling a hit to the target *k* is

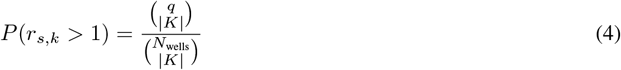

We interpret this probability as the false positive rate for our hit calling metric, or the rate at which we would call a sequence a hit to a given target under the conditions that it lacks specific binding in every well.

For hit calling in *Library 1*, we have *N*_*wells*_ = 296 with |*K*| = 4 replicates per target and choose *q* = 12, such that any sequence can be called a hit to at most 3 unique targets and our theoretical false positive rate is 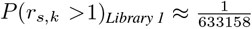 For hit calling in *Library 2*, we have *N*_*wells*_ = 512 with |*K*| = 4 replicates per target and choose *q* = 20, such that any sequence can be called a hit to at most 5 unique targets and our theoretical false positive rate is 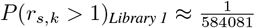. In both library experiments, we include a substantial number of empty wells in addition to wells containing target proteins, to enhance statistical power and rule out plastic-binding hits.

### A.11 Hypothesis Testing for On-Design Hit Rate Significance

For each of the 145 experimentally tested targets, we assess whether designs created specifically for a target achieve a higher hit rate than designs intended for other targets. Let

*D*_*k*_ ≡ {designs against target *k*}, *D*_*k*_*′* ≡ {designs against all targets except *k*}, *H*_*k*_ ≡ {hits against target *k*}.

The set of *on-design hits* is given by *D*_*k*_ ∩ *H*_*k*_, while the set of *off-design hits* is *D*_*k*_*′* ∩ *H*_*k*_.

To formally test whether on-design hit rates are elevated relative to off-design hit rates, we construct a 2 × 2 contingency table for each target *k*:

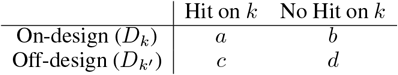

Where

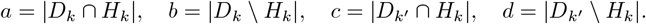

The null hypothesis *H*_0_ posits that the probability of a design hitting target *k* is the same regardless of whether it was generated for *k* or for other targets:

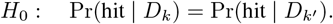

We apply Fisher’s exact test to each contingency table. This test, which is exact even with small sample sizes and varying numbers of designs per target, evaluates whether the observed enrichment of hits in the on-design row (relative to the off-design row) could plausibly arise by chance under *H*_0_. The resulting p-value quantifies the strength of evidence against *H*_0_: small values indicate that the observed difference in hit rates is unlikely under the null, supporting the conclusion that on-design hits are enriched.

## Supplementary Figures and Tables

**Table S1:**
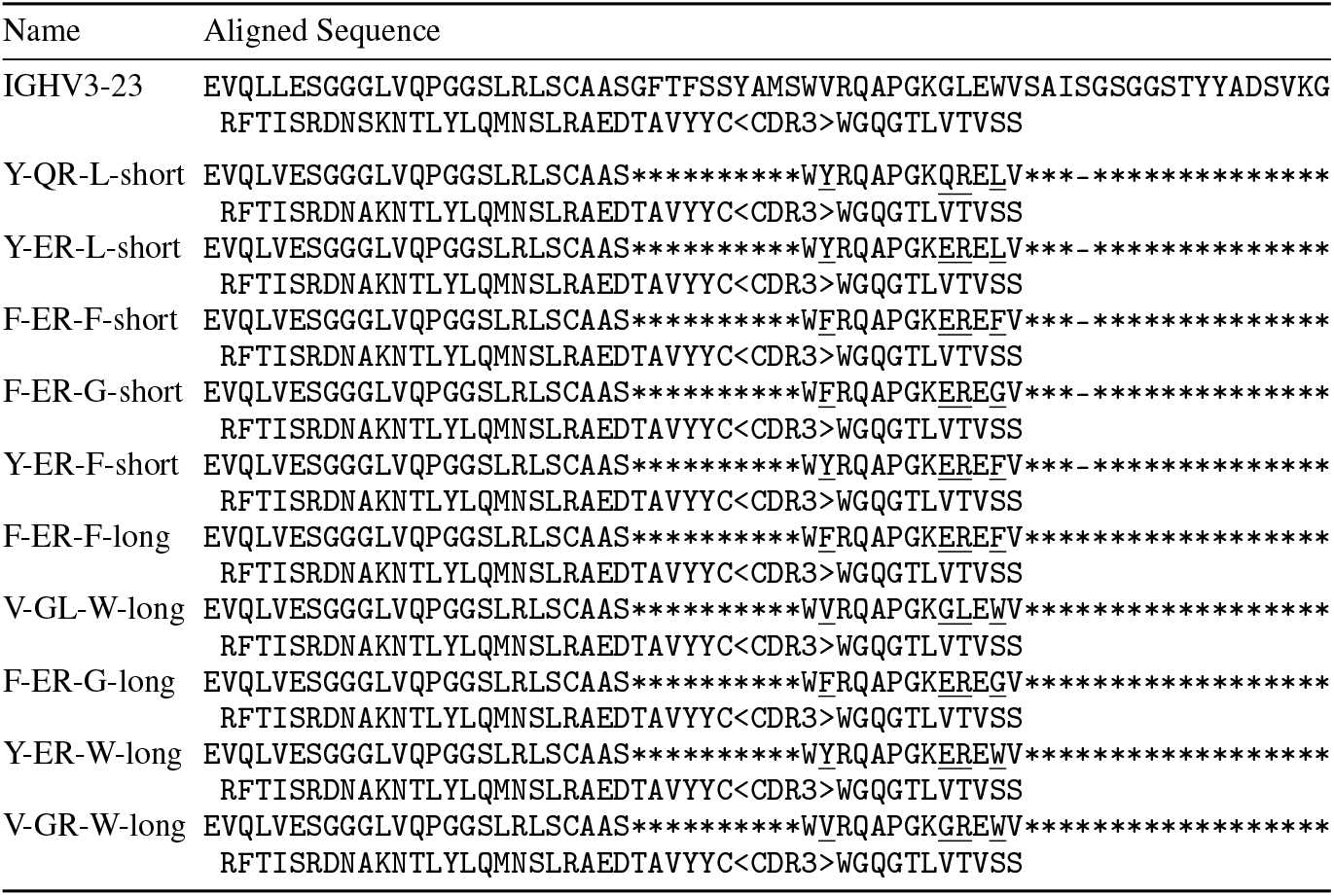
VHH Framework Sequences Based on IGHV3-23. Hallmark residues are underlined. ‘*’ characters indicate positions that are masked and decoded in the process of generation. ‘-’ characters indicate gap positions. We use ‘<CDR3>’ to denote the variable-length CDR3 region. Outside of the variable and hallmark residues, the VHH frameworks differ from IGHV3-23 by the mutations L5V and S74A.

**Figure S1:**
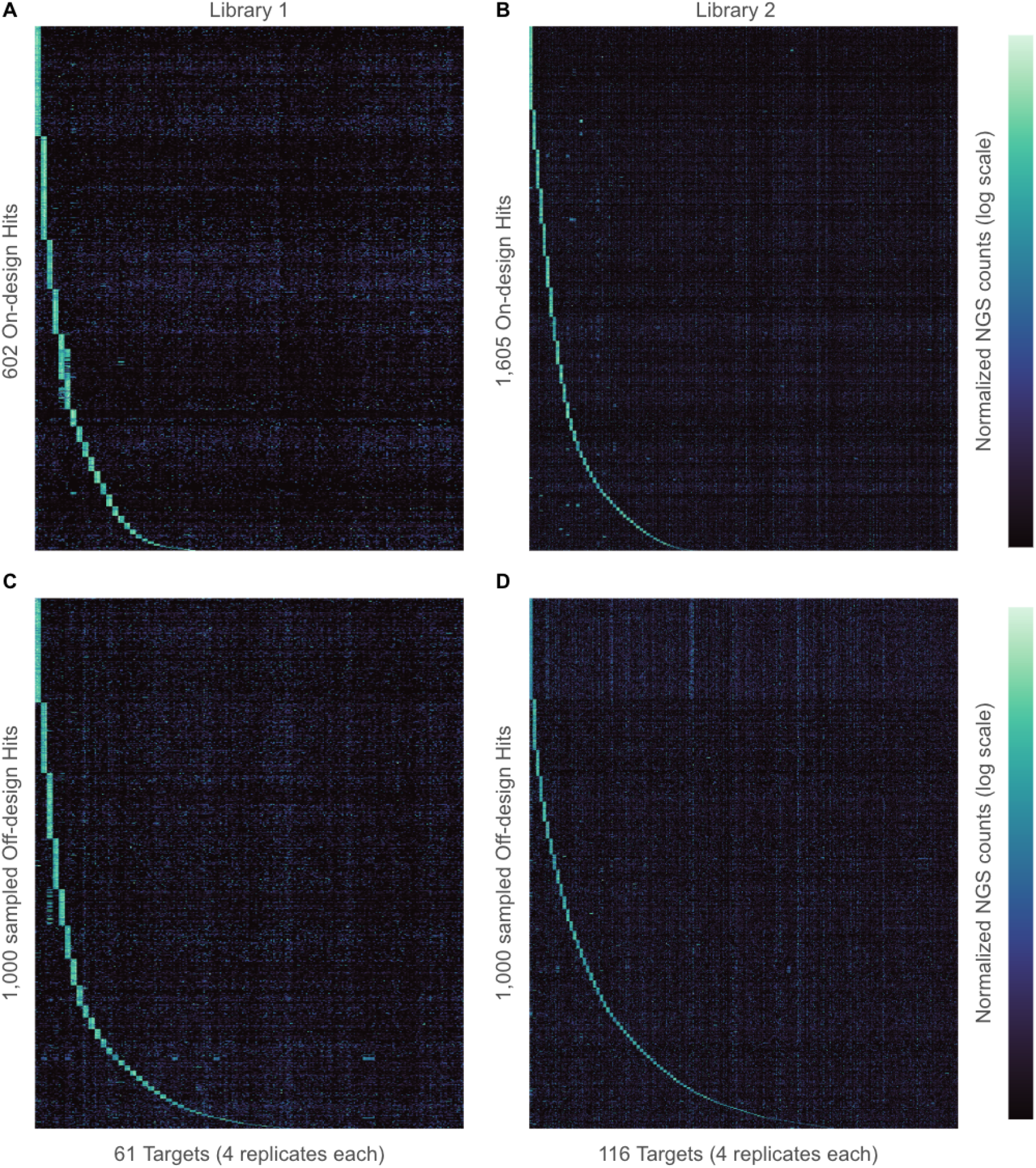
Normalized NGS counts data is shown for sequences called as hits in *Library 1* and *Library 2*. Each row corresponds to a single sequences. Columns correspond to targets, where each target has 4 replicate columns grouped together. Rows and columns are sorted in descending order of number of hits called for visual clarity. **(A)** 602 on-design hits from *Library 1* **(B)** 1,605 on-design hits from *Library 2* **(C)** 1,000 sampled off-design hits from *Library 1* **(D)** 1,000 off-design hits from *Library 2*.

**Figure S2:**
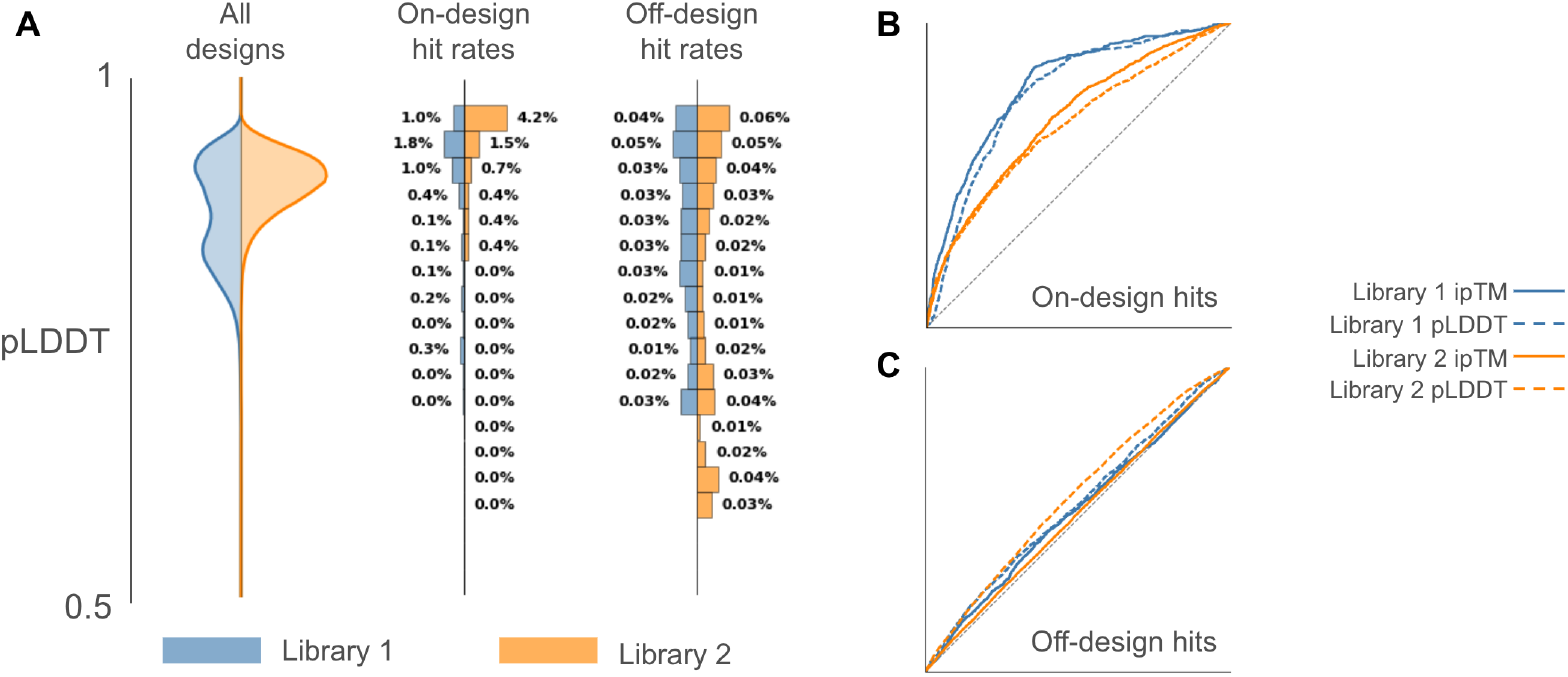
**(A)** Distribution of on- and off-design hit rates binned by pLDDT cutoffs of width 0.025. We filter out bins containing less than 0.1% of each library. Off-design hit rates are normalized by the number of unique targets tested in each library experiment. **(B**,**C)** ROC curves display the discriminatory power of ipTM and pLDDT in each library for filtering to on- and off-design hits.

**Figure S3:**
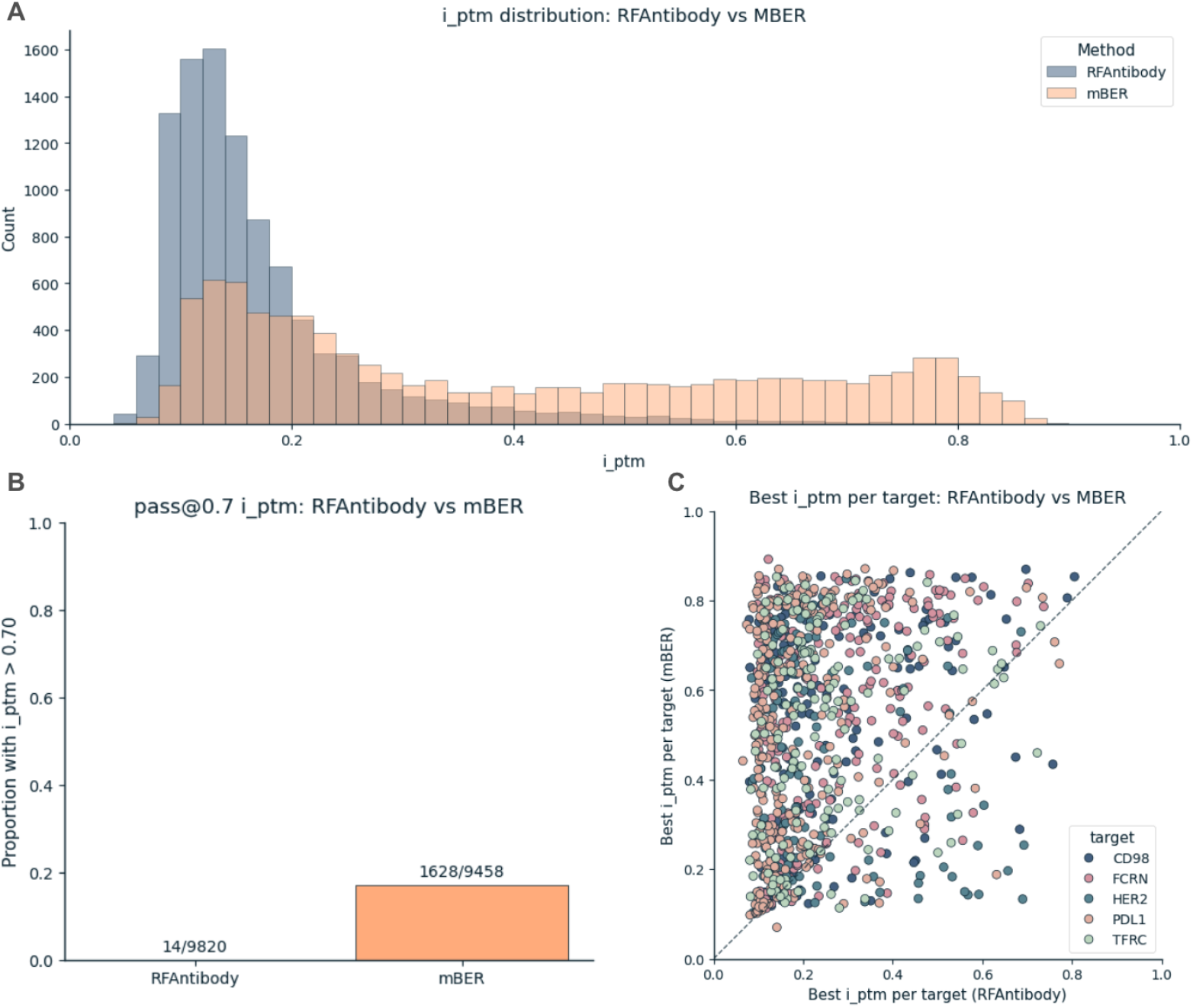
Benchmarking mBER against RFAntibody. **(A)** A histogram of ipTM values for VHH binders generated by RFAntibody and mBER. **(B)** Proportion of binders generated by RFAntibody and mBER passing a cutoff ipTM score of 0.7. **(C)** ipTM values for the best generation from RFAntibody and mBER for matching design configurations. Design configurations are colored by the design target.

**Figure S4:**
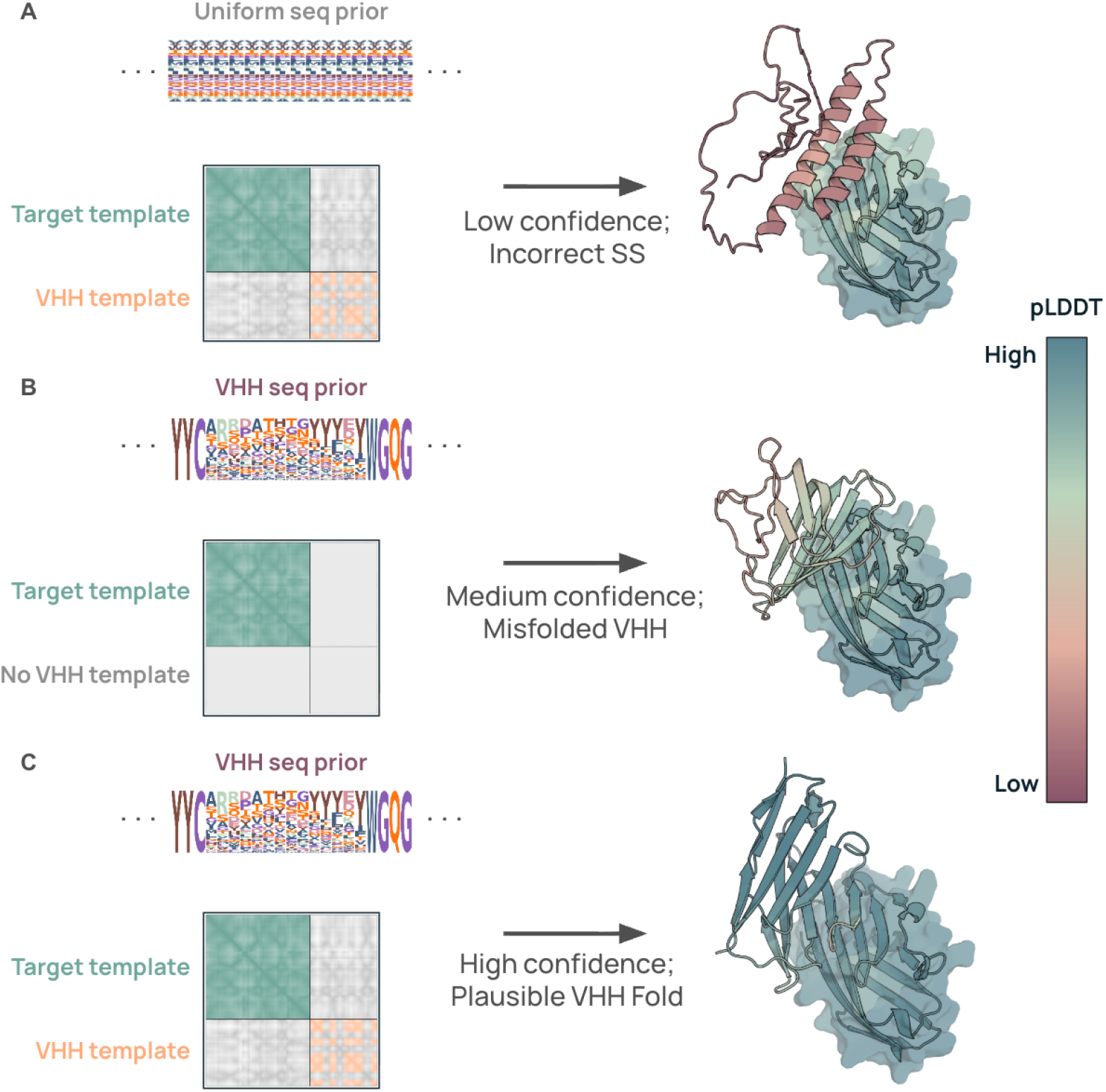
Example binder-target complexes for a successful hotspot on PDL1 when **(A)** a VHH structural template is provided along with the target template to AF2, but with no sequence prior. **(B)** An informative sequence prior is provided to AF2, but the VHH structural template is ablated (target template is still provided). **(C)** Both the sequence prior and VHH structural template are provided to AF2.

## Notes

### Competing Interest Statement

All authors are employees of Manifold Bio.

